# Persistent transmission of shigellosis in England is associated with a recently emerged multi-drug resistant strain of *Shigella sonnei*

**DOI:** 10.1101/798835

**Authors:** Megan Bardsley, Claire Jenkins, Holly D Mitchell, Amy FW Mikhail, Kate S Baker, Kirsty Foster, Gwenda Hughes, Timothy J Dallman

## Abstract

**Background:** Whole genome sequencing has enhanced surveillance and facilitated detailed monitoring of transmission of *Shigella* species in England.

**Methods:** We undertook an epidemiological and phylogenetic analysis of isolates from all cases of shigellosis referred to Public Health England between 2015 to 2018 to explore recent strain characteristics and transmission dynamics of *Shigella* species.

**Results:** Of the 4950 confirmed cases of shigellosis identified during this period, the highest proportion of isolates were *S. sonnei* (54.4%), followed by *S. flexneri* (39.2%), *S. boydii* (4.1%) and *S. dysenteriae* (2.2%). Most cases were adults (82.9%) and male (59.5%), and 34.9% cases reported recent travel outside the UK. Throughout the study period diagnoses of *S. flexneri* and *S. sonnei* were most common in men with no history of recent travel abroad. Species prevalence was not static with cases of *S. flexneri* in men decreasing from 2015-2016, and the number of cases of *S. sonnei* increasing from 2017. Phylogenetic analysis showed this recent increase in *S. sonnei* was attributed to a novel clade exhibiting resistance to ciprofloxacin and azithromycin, which had emerged from a Central Asia sub-lineage.

**Conclusions:** Despite changes in species prevalence, *Shigella* diagnoses in England are persistently the most common in adult males without reported travel history, consistent with sexual transmission amongst men who have sex with men. The trend in increasing ciprofloxacin resistance, in addition to plasmid-mediated azithromycin resistance, in *S. sonnei* is of significant public health concern with respect to transmission of multi-drug resistant gastrointestinal pathogens and the risk of treatment failures.

**Summary:** *Shigella* diagnoses in England are most prevalent amongst men who have sex with men. *S. sonnei* exhibiting resistance to both azithromycin and ciprofloxacin has replaced azithromycin-resistant *S. flexneri* as the most commonly isolated species in this setting.

## Introduction

The four species of the bacterium *Shigella* (*S. sonnei, S. flexneri, S. boydii and S. dysenteriae*) cause dysentery and are transmitted via the faecal-oral route. Shigellosis is the second leading cause of diarrhoeal deaths worldwide, and the highest burden is found in lower-to-middle income countries [1]. In high income countries, shigellosis has historically been associated with either travel-associated infections or self-limited transmission in community risk-groups. However, since being identified as a sexually transmitted infection (STI) in 1974 in San Francisco’s gay community [2, 3], sustained sexual transmission has become an important component of *Shigella* spp. epidemiology. Outbreaks of *S. sonnei* and *S. flexneri* among gay, bisexual and other men who have sex with men (GBMSM) have been reported in urban settings across North America, Europe, Asia and Oceania [4–12].

In England, *Shigella* spp. epidemiology has changed markedly in the past decade; from being a predominantly travel-associated infection, to one where domestically-acquired infections in men account for a large and increasing proportion of diagnoses, with national surveillance data, case questionnaires and outbreak investigations suggesting considerable and sustained sexual transmission within specific sexual networks [4, 13, 14]. Since 2009, overlapping GBMSM-associated epidemics of *S. flexneri* serotype 3a, followed by *S. flexneri* serotype 2a, and more recently *S. sonnei*, have emerged, while case numbers have remained stable in females and travel-associated cases [13].

Whole genome sequencing (WGS) studies have been used to determine factors driving the emergence and transmission of *Shigella* spp. and have demonstrated that certain community outbreaks of shigellosis in the UK belonged to prolonged, global epidemics [5, 15-18]. These analysis have shown that lineages of *S. flexneri* serotypes 3a and 2a have exhibited intercontinental spread via sexual transmission in GBMSM networks, and subsequently acquired multiple antimicrobial resistance (AMR) determinants [5, 17]. Within the *S. sonnei* population, Baker *et al*. (2018) [16] defined four clades associated with transmission in GBMSM communities, all within lineage III the dominant extant lineage both in the UK and globally [19, 20]. A two-year population-level study of all cultured *Shigella* isolates in the state of Victoria, Australia (2016-2018) identified 2 predominant MSM lineages circulating [21]; the *S. flexneri* 2a lineage described above and an MDR *S. sonnei* clade associated with reduced susceptibility to azithromycin, trimethoprim-sulfamethoxazole and ciprofloxacin.

The implementation of WGS in 2015 at Public Health England (PHE) has enhanced surveillance and facilitated the monitoring of transmission of *Shigella* spp.in England [20, 22]. This article uses national surveillance data from 2015 to 2018 to describe the epidemiology of *Shigella* spp. in England in the post-WGS era. We undertook epidemiological and phylogenetic analyses of *Shigella* isolates referred to the GBRU to explore recent characteristics and patterns of transmission of shigellosis in England to better inform the public health response. We examined trends in travel-associated and non-travel associated acquisition by gender to assess evidence for ongoing sexual transmission of *Shigella* spp. among men and report on genomic subtype and antimicrobial resistance changes across the population.

## Methods

### Data collection

The UK Standards for Microbiology Investigation of Faecal Specimens for Enteric Pathogens recommend testing of all faecal specimens for *Shigella* spp. in individuals reporting symptoms of gastrointestinal disease (https://www.gov.uk/government/publications/smi-b-30-investigation-of-faecal-specimens-for-enteric-pathogens). Approximately two-thirds of *Shigella* spp. positive isolates are submitted from diagnostic laboratories to Public Health England’s Gastrointestinal Bacterial Reference Unit (GBRU) for confirmation of species identification and typing, in concordance with varied local practice. Subsequent microbiological typing, including confirmation of species, serotype, and single nucleotide polymorphisms (SNPs) typing, and patient demographic data including, sex, age and recent travel collected from laboratory request forms upon isolate submission are stored in an integrated molecular national surveillance database.

### Data analysis

Microbiological typing data from all isolates of *Shigella* spp. submitted to GBRU between 2015 and 2018 were extracted and analysed. Travel-associated cases were defined as those reporting recent foreign travel to any country (including both endemic high-risk countries and low-risk countries) seven days prior to the onset of symptoms, based on information from laboratory reports. Travel history is captured for between 60-70% of cases of *Shigella* spp.; and therefore, if no travel history is reported it cannot be inferred that the case did not travel. Laboratory surveillance data lacks patient sexual orientation. However, in line with previous work, we calculated gender ratios as an indicator of cases that may be attributed to sexual transmission among GBMSM [13, Mitchell *et al.* Microbial Genomics (accepted)].

### Whole genome sequencing

Since 2015, microbiological typing has been performed at PHE using WGS [20, 22]. DNA was extracted for sequencing on the Illumina HiSeq 2500 instrument. Quality and adapter trimmed Illumina reads aligned to the *S. sonnei* reference genome Ss46 (Genbank accession: NC_007384.1) or the *S. flexneri* serotype 2a strain 2457T (AE014073.1) using BWA MEM v0.7.12 [23]. Single Nucleotide Polymorphisms (SNPs) were identified using GATK v2.6.5 [24] in unified genotyper mode. Core genome positions that had a high-quality SNP (>90% consensus, minimum depth 10x, MQ >= 30) in at least one isolate were extracted and RaxML v8.2.8 [25] used to derive the maximum likelihood phylogeny of the isolates after first removing regions of the genome predicted to have undergone horizontal exchange using Gubbins v2.0.0 [26]. Hierarchical single linkage clustering was performed on the pairwise SNP difference between all isolates at various distance thresholds (Δ250, Δ100, Δ50, Δ25, Δ10, Δ5, Δ0) [27]. Genome-derived serotyping and AMR-conferring genes and SNPs was performed using the *GeneFinder* tool (https://github.com/phe-bioinformatics/gene_finder) [21, 28]. Resistance to streptomycin, tetracycline, the sulphonamides and trimethoprim in *Shigella* species is high, and trends have been consistent for many years [5, 15-17, 28]. We therefore focused our AMR analysis on the presence of genomic makers of resistance to macrolides (specifically *ermB* and *mphA*) and fluroquinolones (specifically mutations in the quinolones resistance determining regions (QRDR) of *gyrA* and *parC*), as the trends in resistance to these clinically relevant classes of antimicrobials fluctuated during the study period (data not shown). FASTQ reads from all sequences in this study can be found at the PHE Bioproject <u>PRJNA315192</u>. For international context, 110 *S. sonnei* sequences isolated from Australian MSMs, previously reported by Ingle *et al.* (2019), were processed as above for phylogenetic comparison.

## Results

First, we analysed the demographic data available, specifically age, sex and travel history linked to the reference laboratory confirmed cases of shigellosis in England for all *Shigella* species and then for each individual species of *Shigella*. We then analysed the WGS data for genomic markers of resistance to azithromycin (*ermB* and *mphA*) and ciprofloxacin (mutations in the QRDR of *gyrA* and *parC*) for all *Shigella* species and for each individual species of *Shigella*. Finally, we investigated the trends in the number of cases and resistance to azithromycin and ciprofloxacin, among *S. flexneri* 2a and *S. flexneri* 3a and the clades of *S. sonnei* previously associated with sexual transmission in men.

### Epidemiology of shigellosis in England

Between 2015 and 2018, there were 4950 reference laboratory confirmed cases of shigellosis, of which the majority were male (2946/4950, 59.5%) and adult aged ≥16 years (4105/4950, 82.9%). In total, 1725/4950 (34.9%) cases reported recent travel outside the UK. When stratified by age, children (aged <16 years) (262/836, 31.3%) reported significantly less recent travel compared to adults (1462/4105 - 35.6%, chi-squared *p* value <0.05). Among adult shigellosis cases, the proportion of males reporting recent travel outside the UK during this period was significantly lower than females (males: 626/2518, 24.9% vs females: 825/1544, 53.4%, chi-squared *p* value <0.001).

During the study period, the number of isolates of *Shigella* spp. referred from non-travel associated adult male cases exceeded cases reporting recent travel at every time point, although marked fluctuations were observed (Figure 1). An increase of adult, male, non-travel-associated cases was first observed in 2013, as previously reported by Simms *et al*. 2015 and attributed to MSM transmission [13] (Figure 1). During the time-frame of this study, adult, male, non-travel cases of shigellosis remained at epidemic levels throughout the first quarter of 2015 before falling by 46% from 2015 to 2016 (674 to 364 cases), but subsequently re-emerged from 2017 onwards.

**Figure 1:**
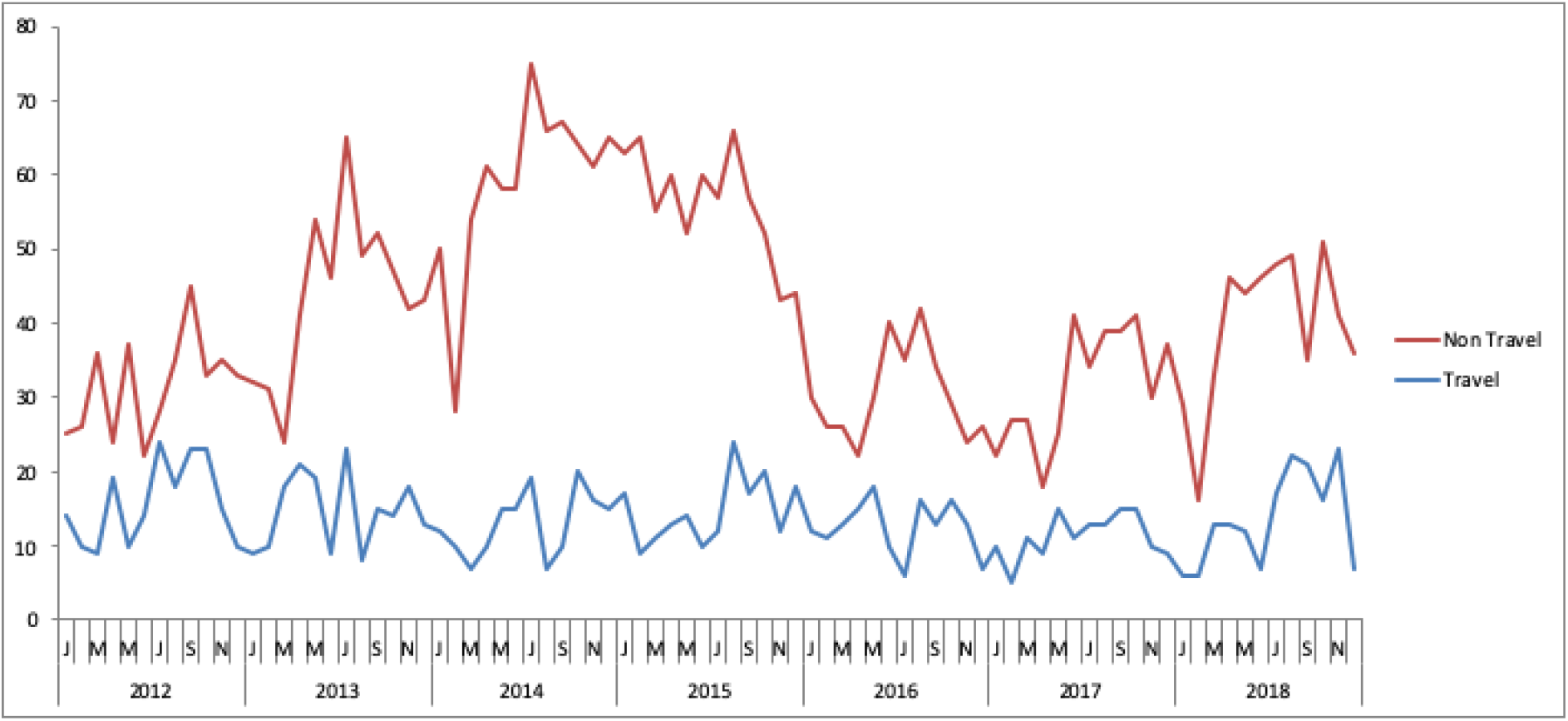
Trends in travel association of *Shigella* spp. diagnoses among men aged ≥16 years in England. Data from Simms *et al*. 2015 [13] analysed prior to this study are included in this figure for context.

### Speciation, serotyping and analysis of patient demography

Of the 4950 isolates of *Shigella* submitted to GBRU between 2015 and 2018, the highest proportion were *S. sonnei* (n = 2695), followed by *S. flexneri* (n = 1943), *S. boydii* (n = 201) and *S. dysenteriae* (n = 111) (Table 1). The most common *S. flexneri* serotypes were 2a (n=1113/1943, 57.2%), 1b (n=254/1943, 13.1%) and 3a (n=206/1943, 10.6%). The most common serotypes of *S. boydii* were serotype 1 (n=24/201, 11.9%), serotype 2 (n=42/201, 20.9%), and serotype 4 (n=23/201, 11.4%), and the most common serotype of *S. dysenteriae* was serotype 2 (n=33/110, 30.0%). All isolates of *S. sonnei* express the same somatic antigen and therefore cannot be serotyped.

**Table 1.** *Shigella* species in England 2015-2108 (n=4950) by age, gender and travel

| Species | Total | Adults<br>(%) | Children<br>(%) | Males<br>(%) | Females<br>(%) | Travel<br>associated (%) | Non-travel<br>associated (%) |
| --- | --- | --- | --- | --- | --- | --- | --- |
| *S. sonnei* | 2695<br>(54.4%) | 2228<br>(82.7) | 467<br>(17.3) | 1472<br>(54.6) | 1223<br>(45.4) | 992 (36.8%) | 1703<br>(63.2) |
| *S. flexneri* | 1943<br>(39.2%) | 1627<br>(83.7) | 316<br>(16.3) | 1342<br>(69.1) | 601<br>(30.9) | 547 (28.2%) | 1396<br>(71.8) |
| *S. boydii* | 201<br>(4.1%) | 164<br>(81.6) | 37<br>(18.4) | 88<br>(43.8) | 113<br>(56.2) | 119 (59.2%) | 82 (40.8%) |
| *S. dysenteriae* | 111<br>(2.2%) | 86<br>(77.5) | 25<br>(22.5) | 44<br>(39.6) | 67<br>(60.4) | 67 (60.4%) | 44<br>(39.6) |

Cases of *S. dysenteriae* (60.4%) or S. *boydii* (59.2%) were significantly more associated with recent foreign travel than cases infected with *S. sonnei* (36.8%) or *S. flexneri* (28.2%) (p-value <0.001) (Table 1). Across all species, a similar proportion of cases were among adults, the lowest proportion was in cases of *S. dysenteriae* (77.5%) and the highest in *S. flexneri* (83.7%). A significantly higher proportion of cases were male for *S. sonnei* (54.6%) and *S. flexneri* (69.1%) compared to *S. boydii* (43.8%) and *S. dysenteriae* (39.6%) (Chi-Squared p-value <0.001) (Table 1).

### Genomic surveillance of markers of antimicrobial resistance to azithromycin and ciprofloxacin

The genome-derived AMR profiles were available for 4057/4950 (81.4%) isolates of *Shigella* in the study. Determinants encoding azithromycin resistance, *ermB* and *mphA*, were most commonly detected in male, non-travel cases of *S. flexneri* and *S. sonnei*, with 45% displaying this resistance genotype compared to <5% non-travel female cases (males reporting travel n=63/660, 9.5%; males, non-travel n=776/1728, 45.0%; females reporting travel n = 5/794, 0.6%; females, non-travel n = 42/875, 4.8%) (Table 2). No azithromycin resistance was detected in *S. boydii* or *S. dysenteriae* isolates. Overall resistance to azithromycin remained stable during the study period, although a downward trend was observed in azithromycin resistance associated with *S. flexneri* isolates from male cases and an upward trend identified in isolates of *S. sonnei* from male cases (Figure 2).

**Table 2.** Presence of AMR determinants encoding resistance to macrolides (*ermB/mphA*) and to fluroquinolones (triple mutations in QRDR) in adult, non-travel associated cases of shigellosis

|  | Male |  |  | Female |  |  |
| --- | --- | --- | --- | --- | --- | --- |
|  | All cases<br>n=1458 | *S. flexneri*<br>n=720 | *S. sonnei*<br>n=738 | Total<br>n=573 | *S. flexneri*<br>n=171 | *S. sonnei*<br>n=402 |
| *ermB/mphA* | 774<br>(53.1%) | 390<br>(54.2%) | 384<br>(52.0%) | 30<br>(5.2%) | 8<br>(4.7%) | 22<br>(5.5%) |
| Triple QRDR<br>mutations | 412<br>(28.3%) | 63<br>(8.8%) | 349<br>(47.3%) | 133<br>(23.2%) | 41<br>(24.0%) | 92<br>(22.8%) |

**Figure 2a.**
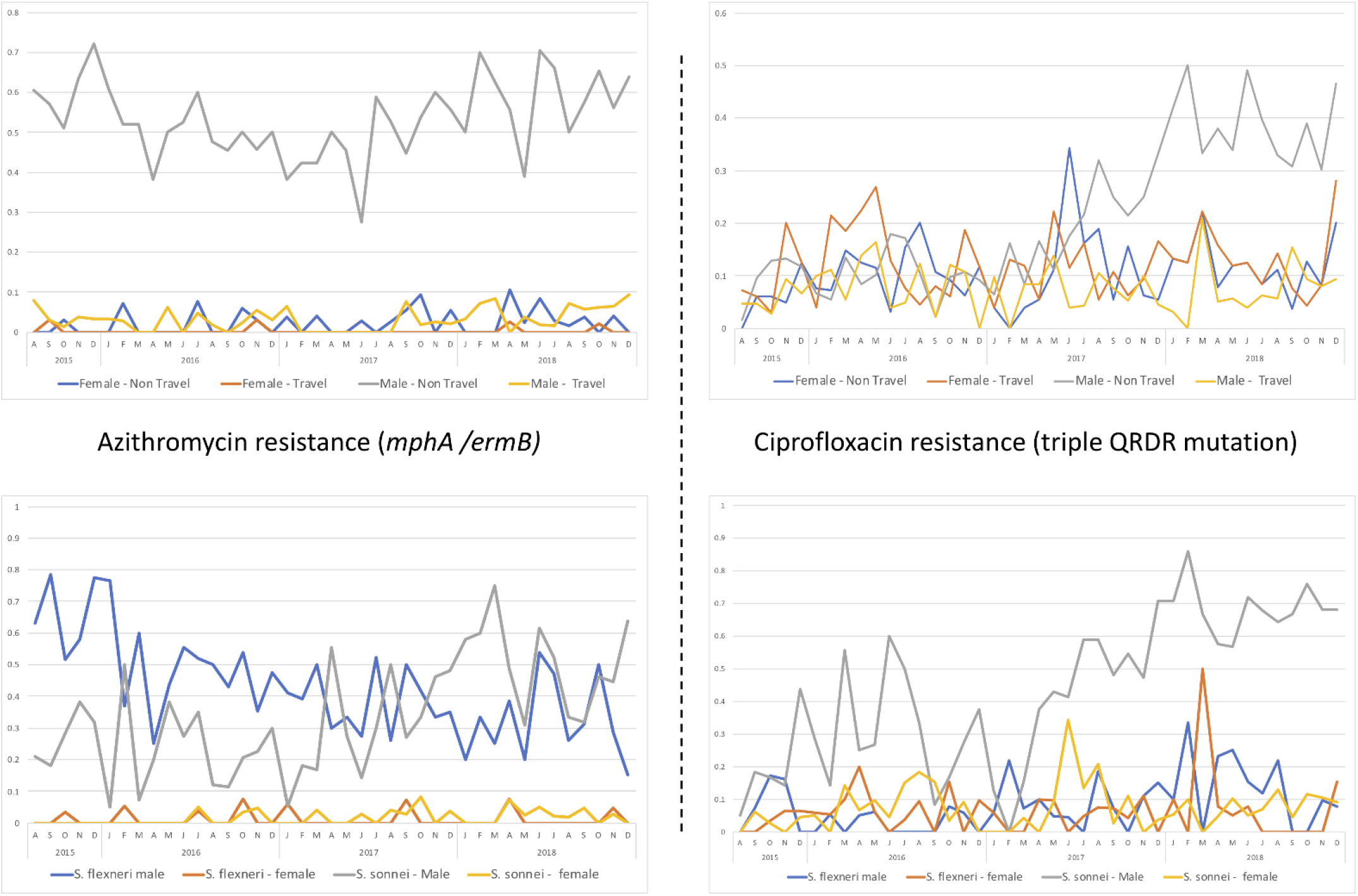
Top boxes illustrate the trends in male and female cases reporting travel outside the UK within 7 days of onset of symptoms and those designated non-travel associated (defined as either reporting no travel or not reporting any travel information) with isolates harbouring genomic markers for resistance to Azithromycin and Ciprofloxacin. Figure 2b. Bottom boxes illustrate the trends in adult male and female cases designated non-travel with isolates of *S. flexneri* or *S. sonnei* harbouring genomic markers for resistance to Azithromycin and Ciprofloxacin.

Triple mutations in the QRDR known to confer resistance to ciprofloxacin were higher in males than females but similar proportions were detected in travel and non-travel cases in both men and woman (males reporting travel n=172/660, 26.1%; males, non-travel n=456/1728, 26.4%; females reporting travel n = 171/794, 21.5%; females, non-travel n = 178/875, 20.3%). Analysis of the available data from adult non-travel cases of *S. flexneri and S. sonnei* showed that triple QRDR mutations were most commonly identified in isolates of *S. sonnei* from males (Table 2). Throughout the study period the proportion of shigellae with ciprofloxacin-resistance conferring QRDR mutations was stable in females and males reporting travel. However, a proportional increase was identified in the male non-travel associated cases. Stratification of these data by *Shigella* species showed that this trend was caused by an increase in the number of isolates of *S. sonnei* (Figure 4). By 2018, 80% of adult male non-travel cases had triple QRDR mutations.

*Phylogenetic trends in S. sonnei* and *S. flexneri* serotype 2a and 3a MSM-associated clades Among all adult cases without recent travel history, *S. flexneri* 2a and *S. flexneri* 3a (previously-implicated in MSM-associated shigellosis epidemics) accounted for 32.3% (1041/3225) of all laboratory-confirmed infections between 2015 and 2018. The age and sex distribution for cases of *S. flexneri* 2a and *S. flexneri* 3a were strongly associated with males (82.3% -860/1041) over the age of 16 years (91.5% - 787/860). The fall in the laboratory-confirmed infections of *S. flexneri* 3a in adult males with no travel history first reported in 2014 has continued throughout this study period (51 cases in 2015 to 17 in 2017, and 15 in 2018) and the number of isolates submitted nationally are now at pre-epidemic levels (Figure 3 and Supplementary figure). Diagnoses of *S. flexneri* 2a in men have also declined, with a 71.4% fall during this study period; from the peak in 2015 (297 cases) cases have fallen annually to 144 cases in 2016, 140 cases in 2017 and just 85 cases in 2018 (Figure 3 and Supplementary figure). Diagnoses from women during this period remained low, below 12 cases in each quarter (Table 1, Figure 2).

**Figure 3.**
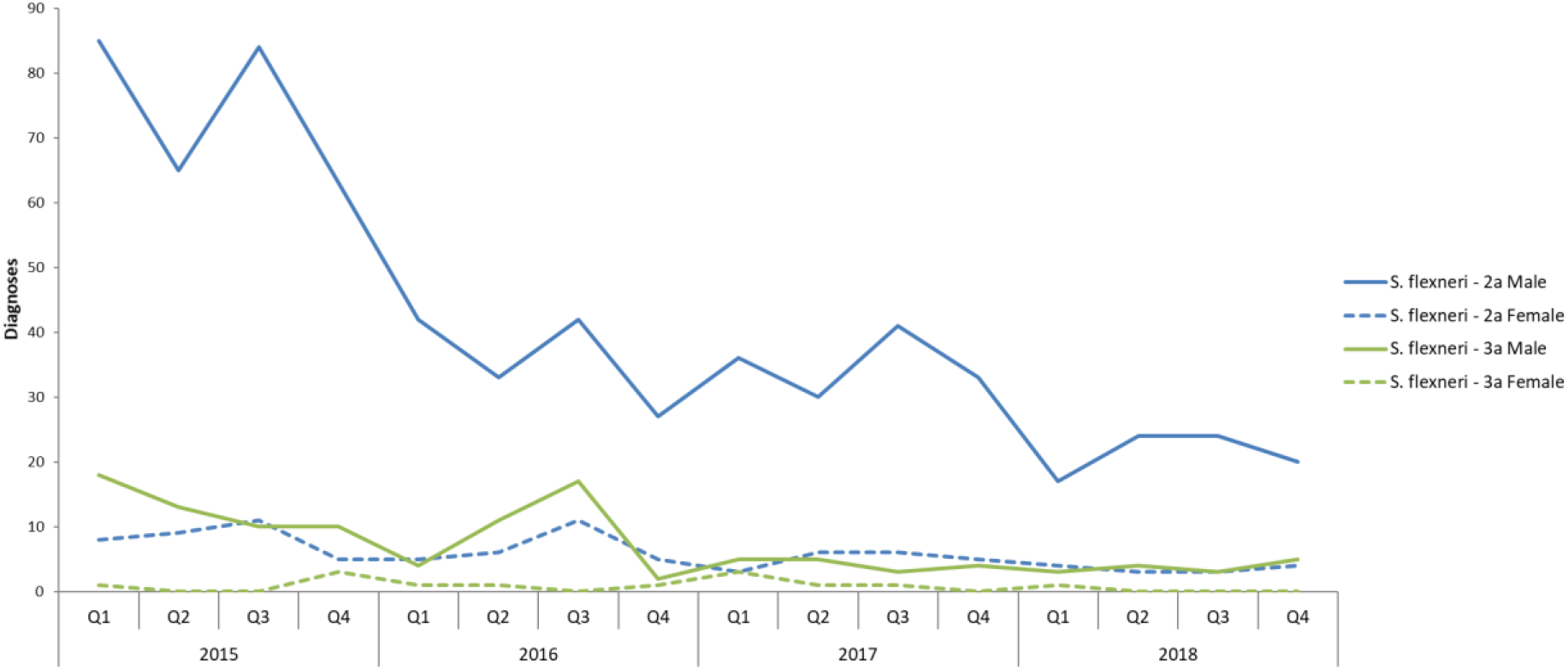
Trends in number of cases of *S. flexneri* 2a and 3a in adult males and females

Phylodynamic analysis of the genome sequences of *S. flexneri* 2a and 3a MSM-associated clades across the two epidemics show that the decrease in case numbers described above, is reflected in decreased sample diversity over time. Bayesian skyline plots reveal the effective population size of *S*. *flexneri* 3a has decreased to pre-epidemic levels, whereas for *S. flexneri* 2a the effective population size is a log-fold change less than the height of the epidemic in 2015 although still above pre-epidemic levels (Supplementary figure).

*S. sonnei*, also previously implicated in MSM transmission, accounted over half (52.8% - 1703/3225) of adult cases without recent travel history between 2015 and 2018. The age and sex distribution for cases of *S. sonnei* was also associated with males (61.8% - 1052/1702) over the age of 16 years (85.9% - 904/1052). Non-travel associated cases of *S. sonnei* in men reduced from 2015-2016, in parallel with *S. flexneri* 2a, but increased again from mid-2017 with the highest number of diagnoses in the study period reported in Q2-Q4 2018 (Figure 4). Female cases have remained relatively stable, except for a notable decline in the first quarter of 2018 and subsequent peak in the third quarter of 2018.

**Figure 4.**
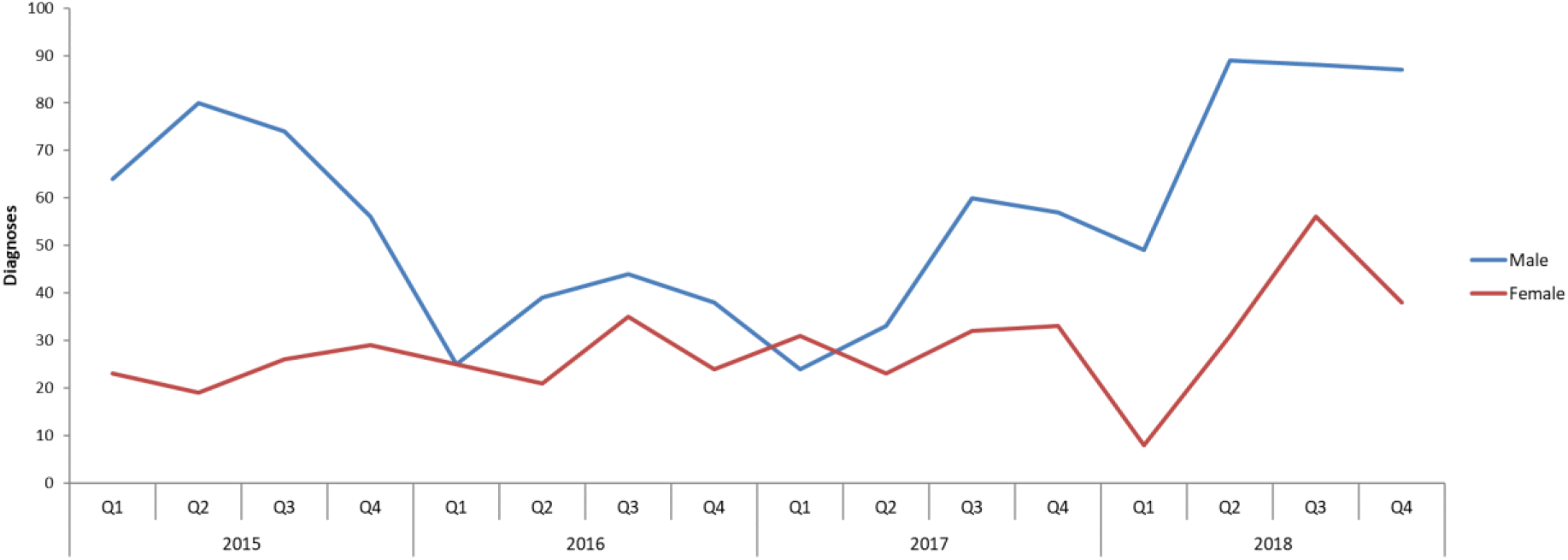
Non-travel associated diagnoses of *Shigella sonnei* among male cases aged ≥16 years.

Phylogenetic analysis was performed to determine whether the four clades of *S. sonnei* previously determined to be circulating in the UK GBMSM population between 2008 and 2014 (*S. sonnei* clades 1-4 in 2018) were contributing to the recent rise in suspect MSM-associated cases of *S. sonnei* (Figure 5). Of the four clades, only clades 2 and 4 contributed cases during the study period, and this was at a stable rate, with both clades having a median of two cases per month. Isolates in clade 2 and clade 4 have both acquired an IncFII plasmid (pKSR100) harboring macrolide resistance genes (*mphA* and *ermB)* on multiple occasions.

**Figure 5a.**
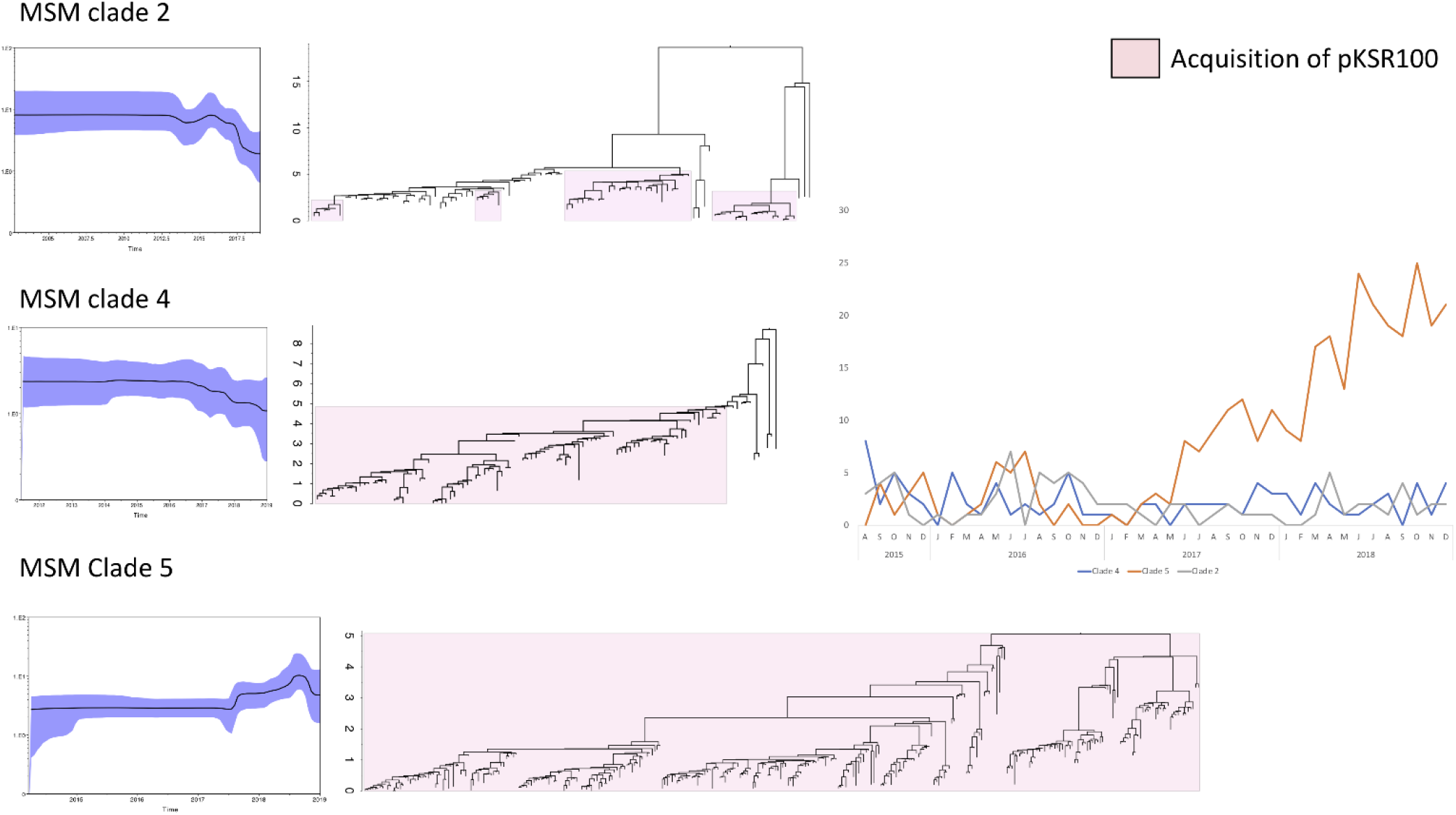
Phylodynamic analysis of *S. sonnei* clades 2, 4 and 5 associated with MSM transmission. Maximum clade credibility trees are presented with Bayesian skyline plots showing the temporal changes of the effective population size of each clade. Figure 5b. Frequency of cases clustering within *S. sonnei* clades 2, 4 and 5 associated with MSM transmission.

In addition, Clade 2 isolates all harbor a single QRDR mutation known to result in reduced susceptibility to ciprofloxacin whereas clade 4 have no QRDR mutations. Phylodynamics analysis suggests that both clades 2 and 4 have a declining effective population size, consistent with the epidemiological evidence of stable case rates attributable to these lineages, suggesting that these clades are not contributing to the recent increase in transmission. The recent increase in cases in this community were attributed to a novel clade, hereby designated Clade 5. Like Clade 1, Clade 5 has emerged from the Central Asia III sub-lineage. Isolates from both Clade 1 and Clade 5 exhibited resistance to ciprofloxacin mediated by triple mutations (gyrA S83L D87G, parC S80I) in the QRDR and both have acquired pKSR100, resulting in macrolide resistance. Phylodynamic analysis suggests Clade 5 originated around 2015 (95% HPD 2014.2 – 2012.7) and the increased isolation of clade 5 *S. sonnei* in male cases is reflected in an increase effective population size of this clade since 2017 (Figure 5).

To place these findings in international context, 110 *S. sonnei* genome sequences obtained from Australian MSMs previously reported by Ingle *et al*. (2019) were analysed to determine which, if any, of the clades identified in this study were contemporaneously circulating globally. Clade 3 isolates accounted for 88% (97/110) of the Australian MSM *S. sonnei* isolates, 6% (7/110) were part of clade 5 and the remaining 6 *S. sonnei* isolates were in other parts of the *S. sonnei* population not previously implicated in MSM transmission. Of the Australian Clade 5 isolates; one was from 2016, four were from 2017 and two were from 2018 with none reporting travel outside of Australia. All 7 isolates contained triple mutations in QRDR and carried the pKRS100 plasmid encoding resistance to azithromycin as reported in the English Clade 5 isolates.

## Discussion

In this study we present evidence that the previous GBMSM-associated epidemics of *S. flexneri* serotypes 2a and 3a have abated, whilst in contrast, non-travel associated transmission of *S. sonnei* among males has intensified since mid-2017 and is associated with the recent emergence of a novel, specific multi-drug resistant clone, Clade 5. Despite changes in species prevalence, *Shigella* spp. diagnoses remain largely restricted to males without reported travel history and concentrated in urban settings, consistent with continued sexual transmission among GBMSM. As with previously observed Shigellosis epidemics associated with MSM transmission, there is evidence of global dissemination of Clade 5 strains between continents. Furthermore, Clade 5 strains have an enhanced ciprofloxacin resistance profile (triple QRDR mutations) compared to other MSM associated lineages circulating and such its emergence represents a significant public health concern.

Research has shown that proactive campaigns through targeted social media and leaflets in sexual health clinics have failed to raise awareness of shigellosis among GBMSM [29]. Despite this, the number of cases of *S. sonnei* and *S. flexneri* 2a diagnoses followed similar trends of decline between August 2015 to August 2017. Whilst *S. flexneri* 2a and 3a continued to fall, we report an emerging epidemic of *S. sonnei* among men in England without travel history from August 2017 onwards*. S. sonnei* is now the most prevalent endemically-acquired species among this population by orders of magnitude. It is possible that changes to the dominant sub-type reflect levels of herd immunity [30], and that new sub-types have the potential to enter and spread within the GBMSM population under the right conditions. Continued monitoring of typing data will provide insight into the future trends in sub-type prevalence and potential strain replacement events.

Studies of ultraorthodox Jewish communities in Israel and abroad [31] have shown that in endemic regions the incidence of *S. sonnei* shigellosis follows a cyclic pattern with epidemics every 2 years. The timing of these cyclical epidemics of *S. sonnei* shigellosis are attributed to the waning rate of natural immunity to the organism [30, 31]. It is likely that a similar phenomenon occurs among GBMSM; natural exposure to *Shigella* spp. through intensive shigellosis epidemics in the GBMSM community may also increase immunity levels in individuals belonging to sexual networks. Herd immunity may be sufficient to temporarily reduce the circulation of *Shigella* species in these networks and prevent epidemics being sustained in subsequent years. However, eventual waning levels of antibodies could lead to a decrease in the herd immunity below a critical level, and thus enable renewed epidemic transmission of shigellosis in MSM. This phenomenon may also facilitate strain replacement events from strains with different immunogenic properties or fitness profiles. Given the indication here that we should anticipate repeated epidemics of strains imported from endemic areas, future studies to develop a predictive framework from genomic information would be beneficial.

Given the ever-increasing reports of AMR in *Shigella* spp. and evidence that strains harbouring mobile resistance-conferring plasmids can spread intercontinentally through sexual transmission [5], therapy guided according to susceptibility testing should be a priority. Previous studies have discussed the role of AMR in driving transmission of shigellosis in the MSM community, and azithromycin resistance is regarded as a marker for isolates associated with sexual transmission among MSM [5, 17]. During the peak of MSM transmission in 2014, only one of the four *Shigella sonnei* clades (clades 1-4) associated with MSM transmission was resistant to ciprofloxacin [16]. Of concern in this study is that the majority of suspect MSM-associated cases are now resistant to both azithromycin and ciprofloxacin. The Clade 5 strain causing the majority of cases in 2017-18 emerged from the same travel-associated clade as clade 1. Given this repeated epidemiological occurrence, it is likely that resistance to ciprofloxacin has driven, or at least contributed to, the emergence and dissemination of clade 5.

Current WHO guidelines recommend the use of fluoroquinolones as the first-line treatment for shigellosis. The increasing trend in resistance to fluroquinolones observed in *S. sonnei* is a public health concern with respect to both the increased likelihood of treatment failures, and to transmission of fluoroquinolone-resistant strains of *S. sonnei* to the wider community. Evidence of transmission of *S. flexneri* previously associated with GBMSM transmission spreading to the wider community has been described [32]. Data on epidemiologically-dominant AMR profiles in *Shigella* spp. would be a beneficial resource to be shared with clinicians to raise awareness and direct appropriate clinical management [33].

Whilst WGS provides highly-detailed and informative information on genetically-linked clusters [32], a lack of patient exposure data, including sexual behavioural data, congregational settings, food and travel history linked to isolates within these clusters hinders the accurate determination of the impact of sexual transmission in shigellosis epidemiology and the most appropriate public health intervention. Collecting enhanced surveillance data for cases of shigellosis is warranted in order to help distinguish between MSM and other forms of transmission requiring different public health actions.

Given the diversity of clinical settings in which shigellosis-affected MSM may present, it is important to consider awareness among clinicians. Healthcare professionals should be aware of the importance of sensitively obtaining sexual history and provide advice about the risk of sexual transmission and the need to avoid sexual activity for at least one week after symptoms cease, and how to prevent onward transmission through sexual and non-sexual conatcts. GBMSM with sexually acquired shigellosis are at high risk of STIs and HIV and should be referred to a sexual health clinic for comprehensive HIV and STI testing and partner notification [4]. From a patient perspective, dense sexual networks, diverse sexual practices including chemsex, and STI and/or HIV co-infection may contribute to the ongoing transmission of *Shigella* spp. among GBMSM [32, 33]. Partner notification and referral for appropriate HIV/STI screening should also be considered. Given the increasing diagnoses of other sexually transmitted infections seen among GBMSM, such as gonorrhoea, syphilis and lymphogranuloma venereum, in the United Kingdom in recent years, [34, 35], https://www.gov.uk/government/statistics/sexually-transmitted-infections-stis-annual-data-tables] and the threat of AMR, strengthened surveillance of *Shigella* spp. transmission is warranted.

---

**Supplementary figure.**
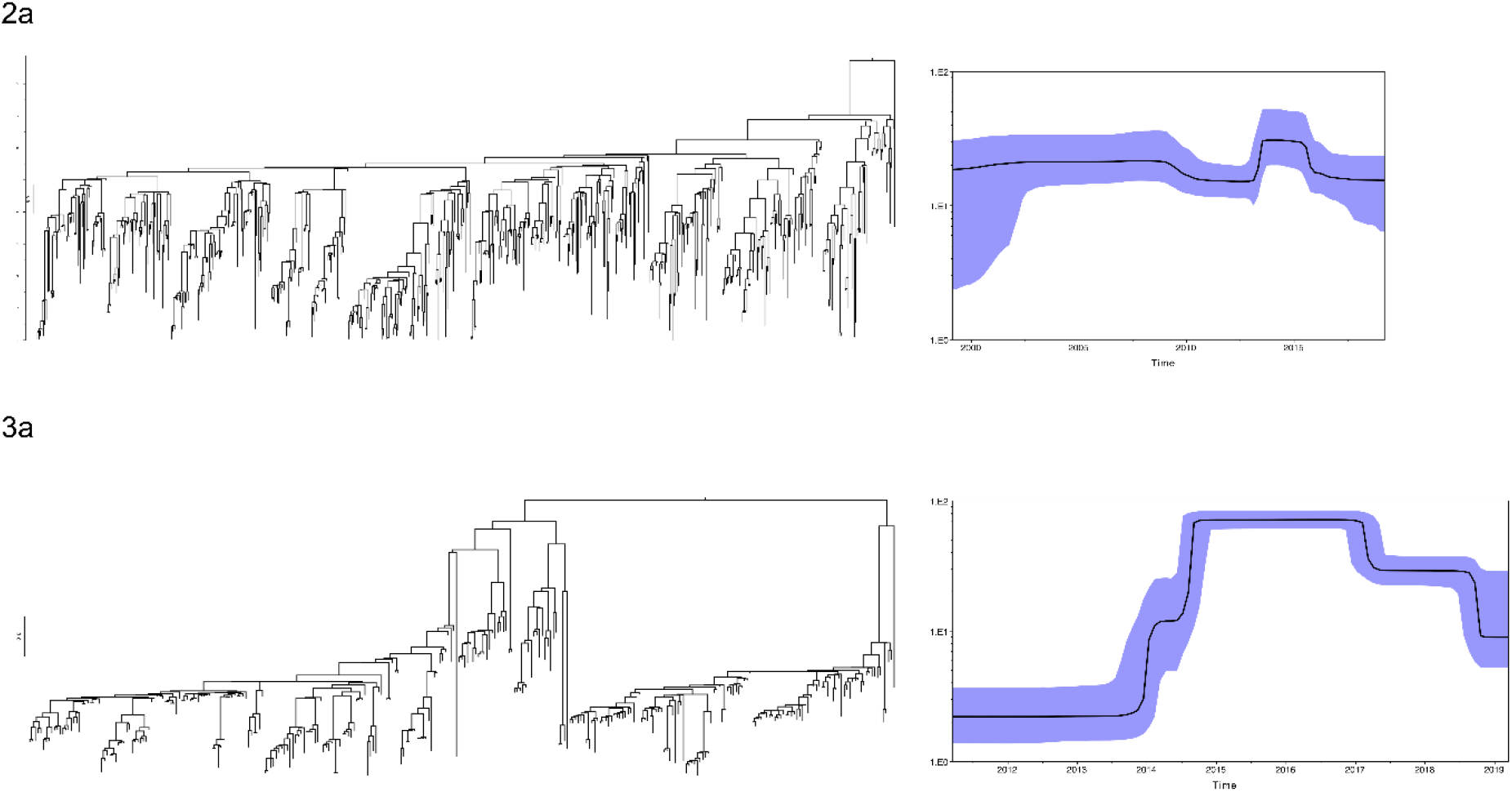
Phylodynamic analysis of *S. flexneri* serotypes 2a and 3a associated with MSM transmission. Maximum clade credibility trees are presented with Bayesian skyline plots showing the temporal changes of the effective population size of each clade.

